# FiNuTyper: an automated deep learning-based platform for simultaneous fiber and nucleus type analysis in human skeletal muscle

**DOI:** 10.1101/2022.12.08.519285

**Authors:** August Lundquist, Enikő Lázár, Nan Sophia Han, Eric B Emanuelsson, Stefan M Reitzner, Mark A Chapman, Kanar Alkass, Henrik Druid, Susanne Petri, Carl Johan Sundberg, Olaf Bergmann

## Abstract

**Summary:** While manual quantification is still considered the gold standard for skeletal muscle histological analysis, it is time-consuming and prone to investigator bias. We assembled an automated image analysis pipeline, FiNuTyper (Fiber and Nucleus Typer), from recently developed deep learning-based image segmentation methods, optimized for unbiased evaluation of fresh and postmortem human skeletal muscle. We validated and utilized SERCA1 and SERCA2 as type-specific myonucleus and myofiber markers. Parameters including myonuclei per fiber, myonuclear domain, central myonuclei per fiber, and grouped myofiber ratio were determined in a fiber type-specific manner, revealing a large degree of gender- and muscle-related heterogeneity. Our platform was also tested on pathological muscle tissue (ALS) and adapted for the detection of other resident cell types (leukocytes, satellite cells, capillary endothelium). In summary, we present an automated image analysis tool for the simultaneous quantification of myofiber and myonuclear types, to characterize the composition of healthy and diseased human skeletal muscle.

**Highlights:**

- A deep learning-based automated platform for skeletal muscle microscopic analysis
- High-fidelity identification and characterization of myonuclei and myofibers
- Validation of SERCA1 and SERCA2 as markers for myofiber and myonuclear subtypes
- Characterization of healthy and pathological human skeletal muscle tissue features
- Adaptations provided for studies on other resident cell types like satellite cells

**eTOC Blurb:** An automated platform for unbiased analysis of skeletal muscle immunohistochemical images, focusing on type-specific myofiber-myonucleus relationships, facilitating high-throughput studies of healthy and diseased tissues.

## Introduction

Histological analysis of muscle structures is crucial for understanding basic muscle physiology and the tissue’s involvement in various pathological conditions. Fundamental research utilizing skeletal muscle histological analysis include studies on exercise, age-related and spaceflight-associated changes in the musculature ^1–8^, but also different forms of dystrophies and neuromuscular disorders, such as amyotrophic lateral sclerosis (ALS) and Parkinson’s disease ^9,10^. Several automated platforms have been developed to investigate skeletal muscle samples in an unbiased fashion, with the potential to analyze them in a high-throughput format. These approaches allow for the measurement of size, shape, and type of myofibers in immunohistochemical images ^11–24^ (Suppl. Table 1). Recently, the ability to measure capillarization and model oxygen consumption for individual myofibers ^12^ and satellite cell identification ^11^ has been added to this toolbox. However, only a few of these automated pipelines allow for simultaneous fiber and myonucleus assignment, based exclusively on the position of the nuclei compared to fiber borders ^16,20^. Routine introduction of myocyte-specific nuclear markers in skeletal muscle histological analysis has been suggested to substantially improve the accuracy and reliability of these types of studies. To date, however, only one such marker has been thoroughly validated in human muscle samples ^25^, and without the direct assignment of myonuclei to a particular fiber type population, its usefulness in studying myofiber-myonucleus relationship is limited ^5,8,26,27^.

Here, we present and validate FiNuTyper, a robust automated platform employing deep learning-based object recognition, for skeletal muscle histological analysis. This novel tool, designed for identifying and quantifying fiber and myonuclear phenotypes, has been optimized for fresh biopsies of healthy and pathological muscle tissue and postmortem samples. In addition, we introduce and validate sarcoplasmic/endoplasmic reticulum calcium ATPase 1 (SERCA1) and 2 (SERCA2) as novel, fiber type-specific myonuclear markers, allowing for simultaneous discrimination of fibers and nuclei of type 2 and type 1 phenotype, respectively.

Using SERCA1- and SERCA2-specific antibodies in combination with cell membrane and nuclear labeling, we established a simple multicolor immunostaining panel for identification and characterization of myocytes and nuclei in human skeletal muscle sections (Fig. 1A). From a set of microscopic images, myofiber and nuclear objects are identified, and subsequently labeled and quantified as type 1 (slow-twitch) or type 2 (fast-twitch), based on their SERCA isoform expression and, in the case of nuclei, also on their vicinity to muscle fibers of the same type. From this information, parameters such as fiber size distribution, number of myonuclei and central myonuclei per fiber, myonuclear domain size (cross-sectional fiber area per myonucleus) and proportion of grouped fibers can be derived separately for both major fiber types (Fig. 1B). In summary, this platform provides a tool for high-throughput histological analysis of the most relevant characteristics of human skeletal muscle in a fiber type-specific manner, facilitating the investigation of skeletal muscle biology in homeostasis and disease.

**Figure 1.**
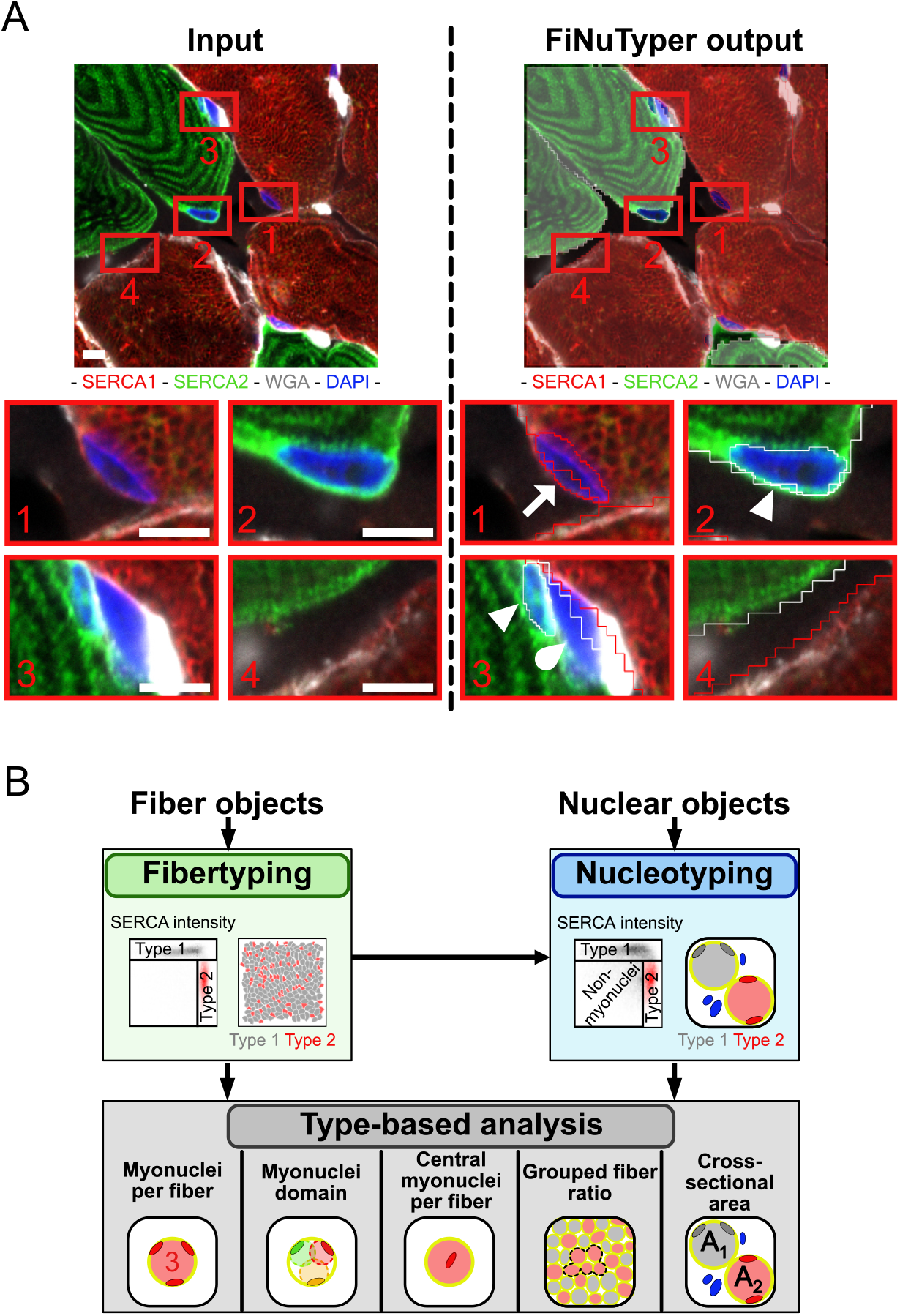
Outline of image processing workflow in FiNuTyper. (A) Input image (left) and graphical output (right) of the analysis performed by FiNuTyper. The areas designated by red rectangles include (1) a SERCA1-positive, type 2 myonucleus (arrow), (2) a SERCA2-positive, type 1 myonucleus (arrowhead), (3) a double negative non-myonucleus (drop) beside a type 1 myonucleus (arrowhead), and (4) an area between fiber borders, not containing any nuclei. The scale bar represents 5 µm. (B) Overview of the image processing pipeline. Fiber objects, generated in Cellpose ^29^ from the WGA fiber border channel are first typed based on SERCA1 and SERCA2 intensity. Nuclear objects, generated in NucleAIzer ^30^ from the DAPI nuclear channel are typed based on SERCA1 and SERCA2 intensity and corresponding fiber type adjacency. The typed objects are co-analyzed to produce measurements of myonucleus per fiber, myonuclear domain, central myonucleus per fiber, grouped fiber ratio, and cross-sectional area, separately for each fiber type.

## Results

### SERCA1 and SERCA 2 are fiber type-specific myonuclear markers

We aimed to establish a simple immunostaining design for the simultaneous, type-specific identification of both myofibers and myonuclei in human skeletal muscle tissue sections, adapted for a classical 4-channel imaging setup available in most fluorescent microscopes. Until now, no myonuclear marker has been described to assign labeled myonuclei to distinct fiber types. Antibodies against slow- and fast-twitch myosin heavy chain isoforms (MyHC1 and MyH2A-2X, respectively), classically used for fiber type determination in human skeletal muscle studies, do not provide a nuclear or perinuclear signal in myofibers. To overcome this challenge, we investigated the potential use of sarcoplasmic reticulum Ca^2+^ pump proteins SERCA1 and SERCA2 as fast- and slow-twitch myofiber and myonuclear markers in human skeletal muscle, respectively. SERCA1 and SERCA2 have been described to show fiber type-specific expression patterns, consistent with the MyHC isoform distribution ^28^. Since the sarcoplasmic reticulum membrane is continuous with the nuclear envelope, we hypothesized that SERCA1 and SERCA2 could be detected in a distinct perinuclear localization besides the sarcoplasmic reticulum membrane, allowing for the identification of both nuclei and fibers of different types in the same section. First, we tested our staining design on transversal sections of postmortem human psoas and pectoralis muscles, combining antibodies against the two SERCA protein isoforms, WGA as cell membrane marker and DAPI as a nuclear stain (Fig. 2A). Most fibers, surrounded by a defined WGA signal, were labeled either by SERCA1 or SERCA2 only, however, we also observed fibers simultaneously expressing both SERCA isoforms, suggesting an intermediate phenotype (Fig. 2A).

**Figure 2.**
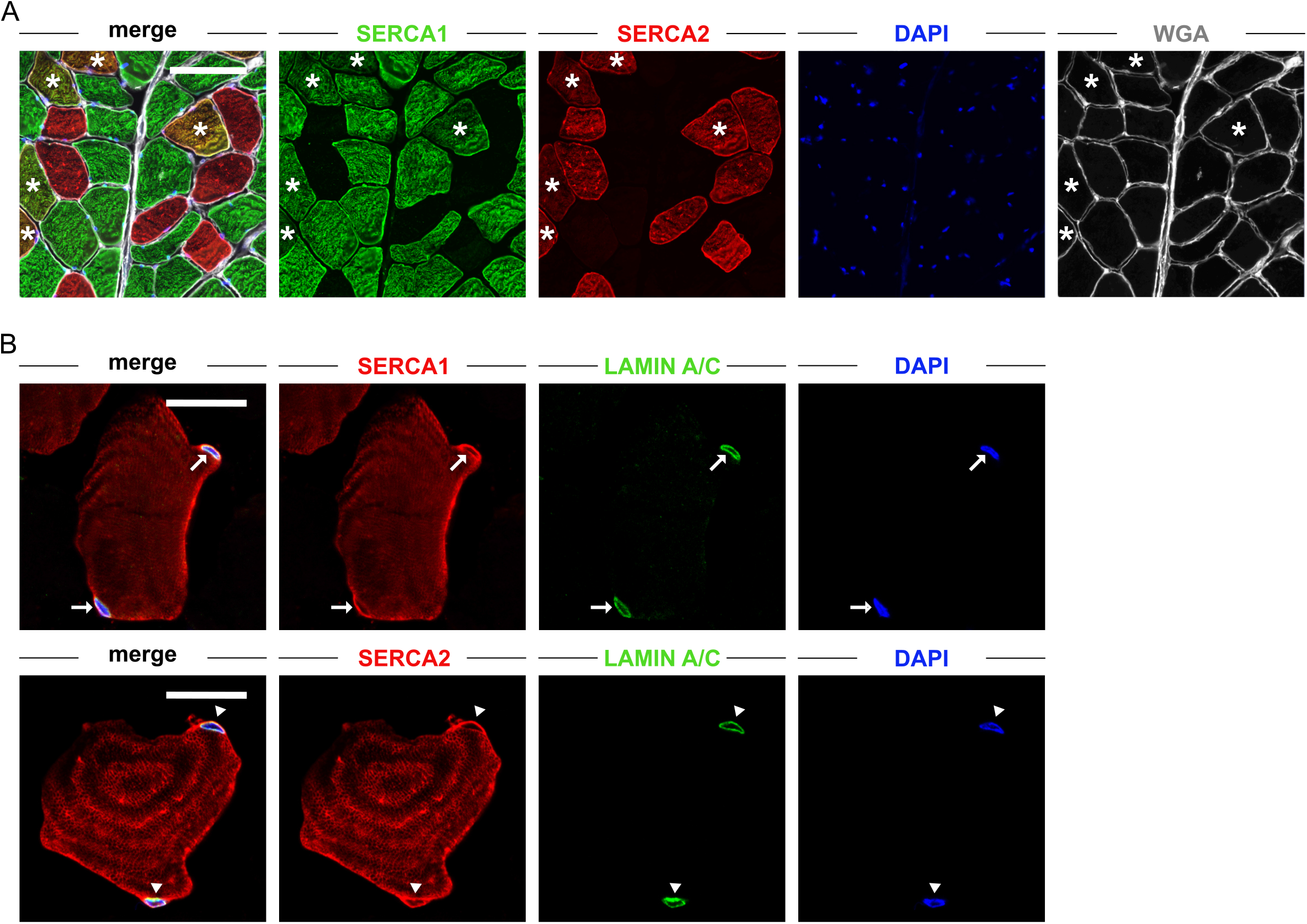
SERCA 1 and SERCA2 show distinct perinuclear localization in complementary myofiber populations. (A) Multicolor immunostaining panel using SERCA1 (green) and SERCA2 (red) as myofiber markers, WGA (white) for fiber border, and DAPI (blue) for nuclear labeling, delineate partially overlapping myofiber populations in transversal sections of human skeletal muscle. The selected area contains a high number of fibers co-expressing both SERCA isoforms (asterisk). The scale bar represents 100 µm. (B) SERCA1 and SERCA2 (red) are detected in a distinct perinuclear structure (besides the sarcoplasmic reticulum of the respective fiber type), consistent with the position of the nuclear envelope, showing close spatial association with the nuclear lamina marker lamin A/C. SERCA1-positive myonuclei are marked by arrows, and SERCA2-positive myonuclei are marked by arrowheads. The scale bar represents 20 µm.

Both SERCA1 and SERCA2 were present in the entire area of the labeled myofibers, in a localization presumably consistent with the position of the sarcoplasmic reticulum (Fig. 2B). Moreover, we detected a distinct perinuclear signal with SERCA1 and SERCA2-specific antibodies in the respective myofibers (Fig. 2B). We confirmed the close spatial association of this signal with the nuclear membrane, by co-staining with lamin A/C- and SERCA1-or SERCA2-specific antibodies (Fig. 2B).

We did not detect any specific SERCA1 or SERCA2 staining in a similar intensity range in non-myocytes outside of the myofiber borders, either in cells dispersed between myocytes (Suppl. Fig. 1A) or more substantial connective tissue borders and vessel walls (Suppl. Fig. 1B), confirming the signal to be highly specific to myofibers and myonuclei. Satellite cells are located between the sarcolemma and basal membrane surrounding myofibers, and as such, could potentially be detected inside the fiber borders defined by the WGA signal. However, we observed no specific SERCA1 or SERCA2 signal around PAX7-labelled satellite cell nuclei (Suppl. Fig. 1C), further confirming their specificity to myonuclei in human skeletal muscle.

To assess the relationship between the expression patterns of type 1-or type 2-specific SERCA and MyHC isoforms, we performed co-staining with combinations of MyHC1-, MyHC2A-2X-, and SERCA1-or SERCA2-specific antibodies (Fig. 3A) on consecutive tissue sections and plotted the mean signal intensities in three parallel channels, measured in individual fibers (Fig. 3B and 3C). The classical fiber type markers MyHC1 and MyHC2A-2X showed complementary expression patterns with little to no overlap, pointing to a low abundance of hybrid fibers in the analyzed samples (Fig. 3A). Most fibers were either colabeled or not labeled by the combination of the type 2-specific SERCA1 and MyHC2A-2X (Fig. 3A, B) and the type 1-specific SERCA2 and MyHC1 antibodies (Fig. 3A, C), corroborating the close association between SERCA and MyHC isoform expression. Moreover, we evaluated the reliability of SERCA1 and SERCA2 as fiber type-specific markers by comparing SERCA-based individual fiber assignment to type 1 and type 2 populations to that based on the MyHC isoform expression pattern, currently considered the gold standard of fiber type analyses (Fig. 3D, E). Based on the dataset collected from postmortem psoas tissue of three subjects covering a wide age range (25, 45 and 73 years) and a stringent gating strategy based on the position of a clearly defined double-positive fiber population, we found both SERCA1 and SERCA2 to be highly sensitive (99.71 ± 0.30% and 100 ± 0.00%, mean ± SD, respectively) and specific (98.77 ± 1.35% and 98.91 ± 1.21%, mean ± SD, respectively) markers of type 2 and type 1 myofibers (Fig. 3F), independent from the age of the individual.

**Figure 3.**
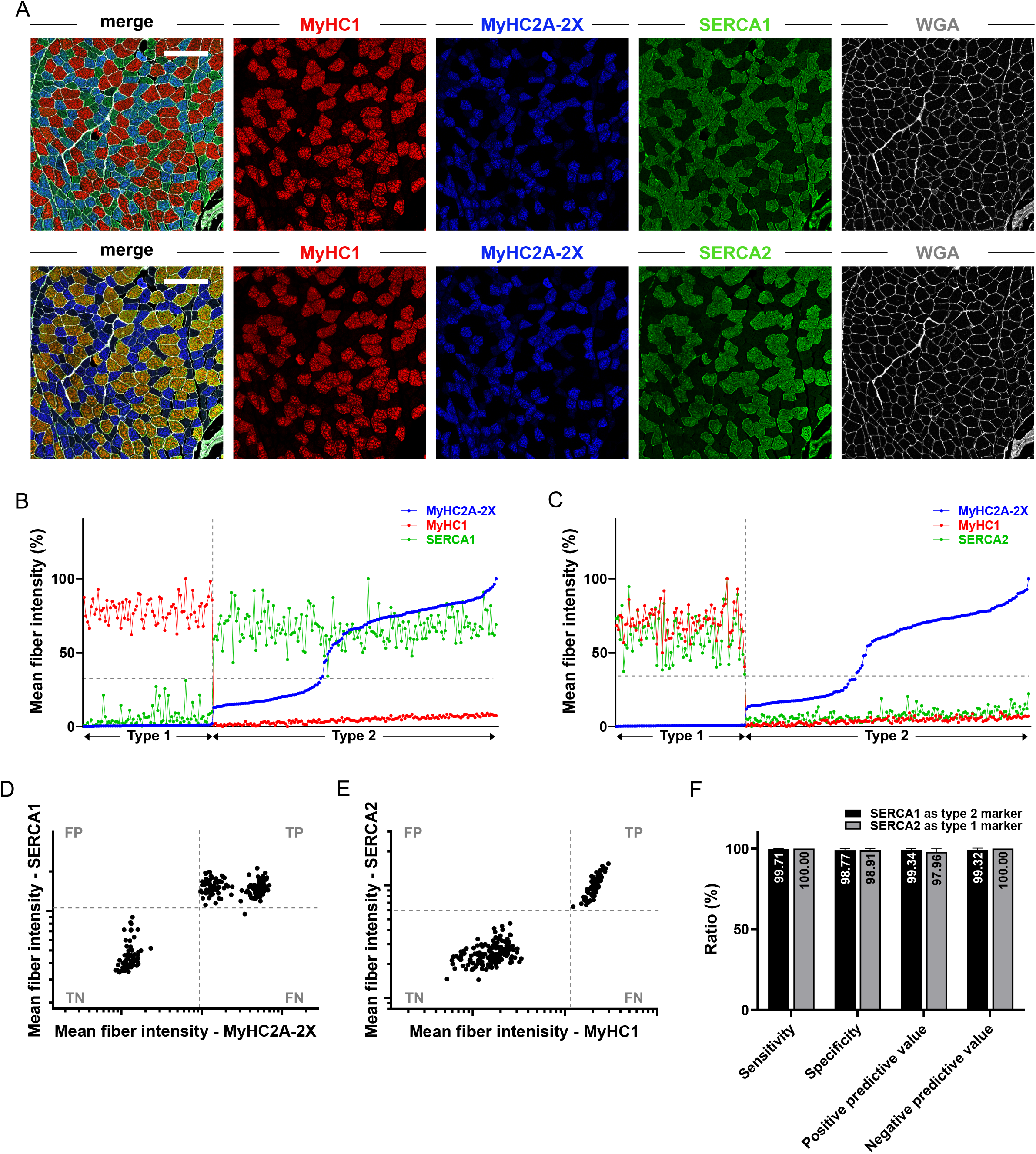
SERCA- and MyHC-based fiber typing approaches provide consistent results in human skeletal muscle. (A) Co-staining of transversal section of human skeletal muscle with antibodies against classical myosin heavy chain markers and the two SERCA isoforms confirm almost complete overlap between SERCA1 (upper panel, green) and MyHC2A-2X (blue), as well as SERCA2 (lower panel, green) with MyHC1 (red) labeling. The scale bar represents 200 µm. (B) Mean signal intensity distribution of SERCA1 (green) and (C) SERCA2 (green) align with type 2 and type 1 fiber identities, assigned based on MyHC2A-2X (blue) and MyHC1 (red) mean signal intensities measured in individual myofibers of the same area (single scans, n=236 fibers). (D) The gating strategy to compare corresponding MyHC- and SERCA isoform-based fiber type assignment for type 2 and (E) type 1 myofibers uses stringent cutoffs, based on the signal intensities of the defined double positive populations. TP: true positives (SERCA^+^-MyHC^+^); FP: false positives (SERCA^+^-MyHC^-^); FN: false negatives: (SERCA^-^-MyHC^+^); true negatives: SERCA^-^-MyHC^-^). (F) Sensitivity, specificity, positive and negative predictive values of SERCA1 as a type 2 (black columns) and SERCA2 as a type 1 (grey columns) myofiber marker (based on n=3 subjects, 3 image scans/subject, mean ± SD).

### FiNuTyper quantifies skeletal muscle microscopy images with high accuracy

After confirming that the sarcoplasmic reticulum- and nuclear envelope-associated SERCA1 and SERCA2 signals allow for simultaneous myofiber- and nucleotyping, we designed a simple immunostaining panel to stain transversal sections of human skeletal muscle, using antibodies against the two SERCA isoforms, combined with fluorescently conjugated WGA as cell membrane marker and DAPI as a nuclear stain (Fig. 1A). Confocal microscopy images, captured from sections processed with this panel, were submitted to FiNuTyper, our automated image analysis pipeline, built on the recently reported image segmentation tools CellPose^29^ and NucleAIzer ^30^. Since the primary output of any fiber-or nucleotyping approach is the number of identified objects and fiber cross-sectional area values, we sought to benchmark our automated tool against manual evaluation, by deriving these parameters from the same image sets and comparing the obtained results.

We assessed the accuracy of FiNuTyper-based fiber (Fig. 4A) and myonuclear identification (Fig. 4B) on frozen sections of fresh vastus lateralis muscle and postmortem psoas major and pectoralis major muscle, by calculating intraclass correlation coefficients (ICC, 95% confidence interval) between the output values of the manual and automated analyses. FiNuTyper displayed an ICC of 0.977 (0.962-0.987) for fiber identification in the postmortem and 0.985 (0.923-0.997) in the fresh biopsy image sets, confirming an excellent agreement between the two independent approaches (Fig. 4A). The high level of accordance between FiNuTyper-based and manual evaluation was also upheld when performing myonuclear identification in the bioptic samples, with an ICC of 0.912 (0.641-0.980) (Fig. 4B). While still showing a good overall correlation, we obtained more variable results in the postmortem dataset with an ICC of 0.790 (0.268-0.918) (Fig. 4B). This is likely due to different postmortem intervals and thus, varying tissue quality of the analyzed samples, which seems to have a more pronounced effect on the reliability of nucleotyping than of fibertyping.

**Figure 4.**
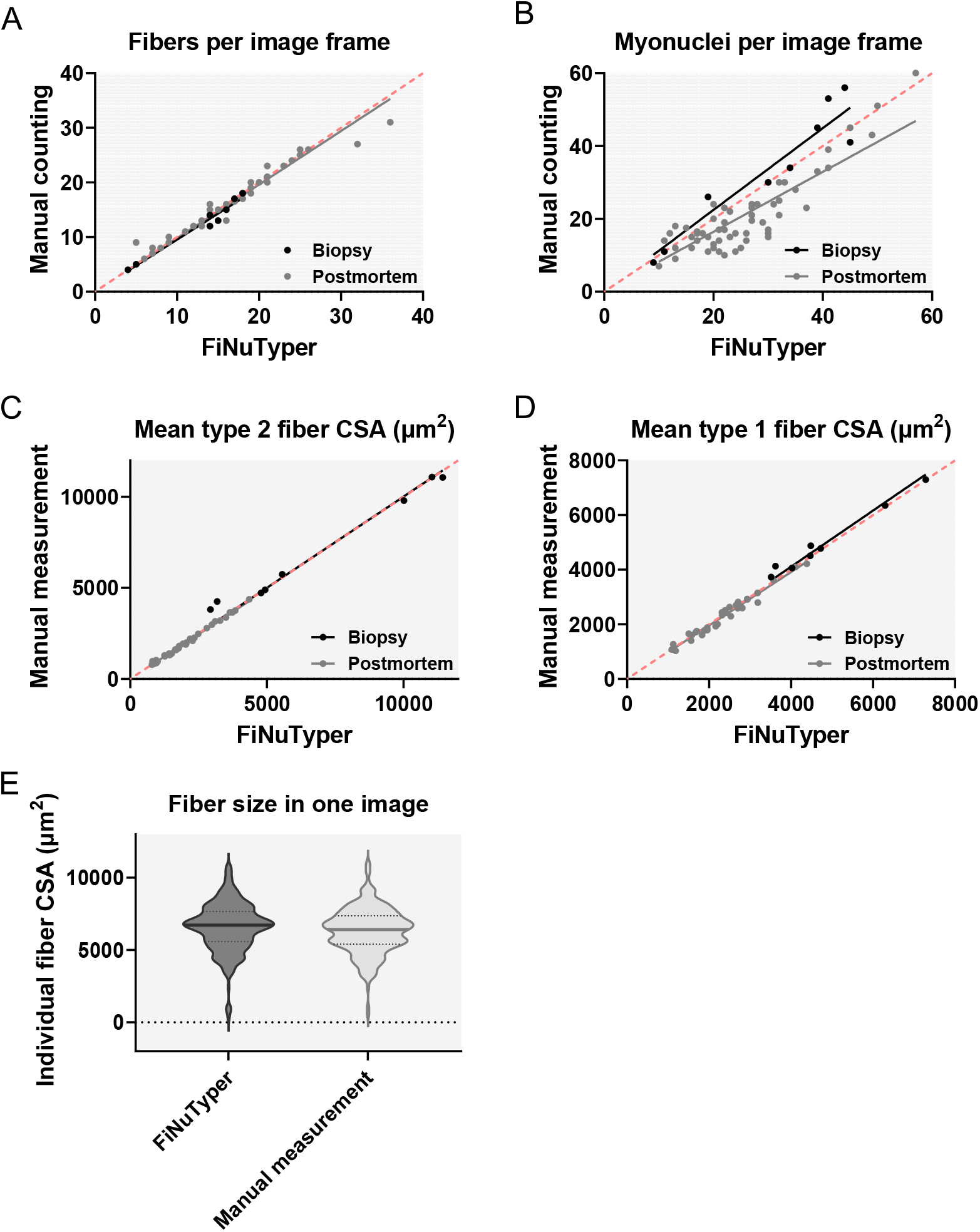
Automated SERCA-based image analysis by FiNuTyper provides results comparable to manual evaluation of human skeletal muscle sections. (A) Comparison of the number of fibers per image frame (ICC=0.985 (0.923-0.997), n=9 for fresh biopsies (black); ICC=0.977 (0.962-0.987), n=57 for postmortem tissue, (grey)); (B) number of myonuclei per image frame (ICC=0.912 (0.641-0.980), n=9 for fresh biopsies (black); ICC=0.790 (0.268-0.918), n=57 for postmortem tissue, (grey)); (C) type 1 fiber cross-sectional area per image frame (ICC=0.988 (0.947-0.998), n=9 for fresh biopsies, (black); ICC=0.997 (0.990-0.999), n=35 for postmortem tissue, (grey)) and (D) type 2 fiber cross-sectional area per image frame (ICC=0.982 (0.855-0.997), n=9 for fresh biopsies (black); ICC=0.985 (0.969-0.993), n=35 for postmortem tissue (grey)), determined manually and generated by the automated approach. Linear regression lines were forced to intersect x,y=0,0. The line of identity is displayed in red. The image frame size was approximately 0.3 × 0.3 mm^2^. ICC: intraclass correlation coefficient, single measure (95% confidence interval). (E) Fiber size distribution in a single image scan (approximately 0.9 × 0.9 μm^2^) based on automated (n=243, dark grey) and manual (n=254, light grey) evaluation.

Since type 1 and type 2 fibers often respond to physiological and pathological challenges by changing their sizes and shapes differently, we decided to validate the automated fiber cross-sectional area (CSA) determination by FiNuTyper (Fig. 4C, D) on a fiber type basis in both postmortem and fresh bioptic samples, against manual measurements. Our analysis revealed a very strong correlation between the manually collected and automatically generated mean CSA values, independent of the fiber type or source of the tissue sample (ICC of type 1 fiber CSA measurements in postmortem tissue: 0.997 (0.990-0.999) and in bioptic tissue: 0.988 (0.947-0.998); ICC of type 2 fiber CSA measurements in postmortem tissue: 0.985 (0.969-0.993) and in bioptic tissue: 0.982 (0.855-0.997)). We performed Bland-Altman analysis on the validation dataset and displayed the results along with the 95% limit of agreement values (Suppl. Fig. 3A-D). We also evaluated the cross-sectional area at the level of individual fibers within a single image scan and found a highly similar fiber size distribution between the manual and automated output (Fig. 4E), further confirming the reliability of CSA measurements performed by FiNuTyper.

### Gender, muscle, and fiber type determine myocyte and myonuclear characteristics

To demonstrate the advantages of FiNuTyper in the analysis of larger image sets, we processed paired samples of postmortem psoas major and pectoralis major muscles of five male and five female individuals, deceased between 44 and 55 years of age, and submitted three image scans (9 stitched frames, approx. 0.9 × 0.9 mm^2^) taken from distinct areas of each muscle sample to the automated pipeline. In total, we identified and analyzed approximately 15600 (780.5 ± 70.7 per muscle sample, mean ± SD) muscle fibers and 12500 (625.4 ± 46.3 per muscle sample, mean ± SD) myonuclei in the processed image scans.

In results compiled from both genders, we found type 1 fibers in significantly higher proportion in the psoas than in the corresponding pectoralis samples (Fig. 5A, p=0.0079), corroborating the postural function of the psoas muscle in contrast to the more dynamic movement profile supported by the pectoralis muscle. The fiber type-specific difference between the two tissues, however, disappeared on the myonuclear level (Fig. 5B). This suggests that type 1 fibers on average contribute more to the myonuclear pool than type 2 fibers, explaining the discrepancy between fiber and myonuclear ratios in the psoas and pectoralis muscles.

**Figure 5.**
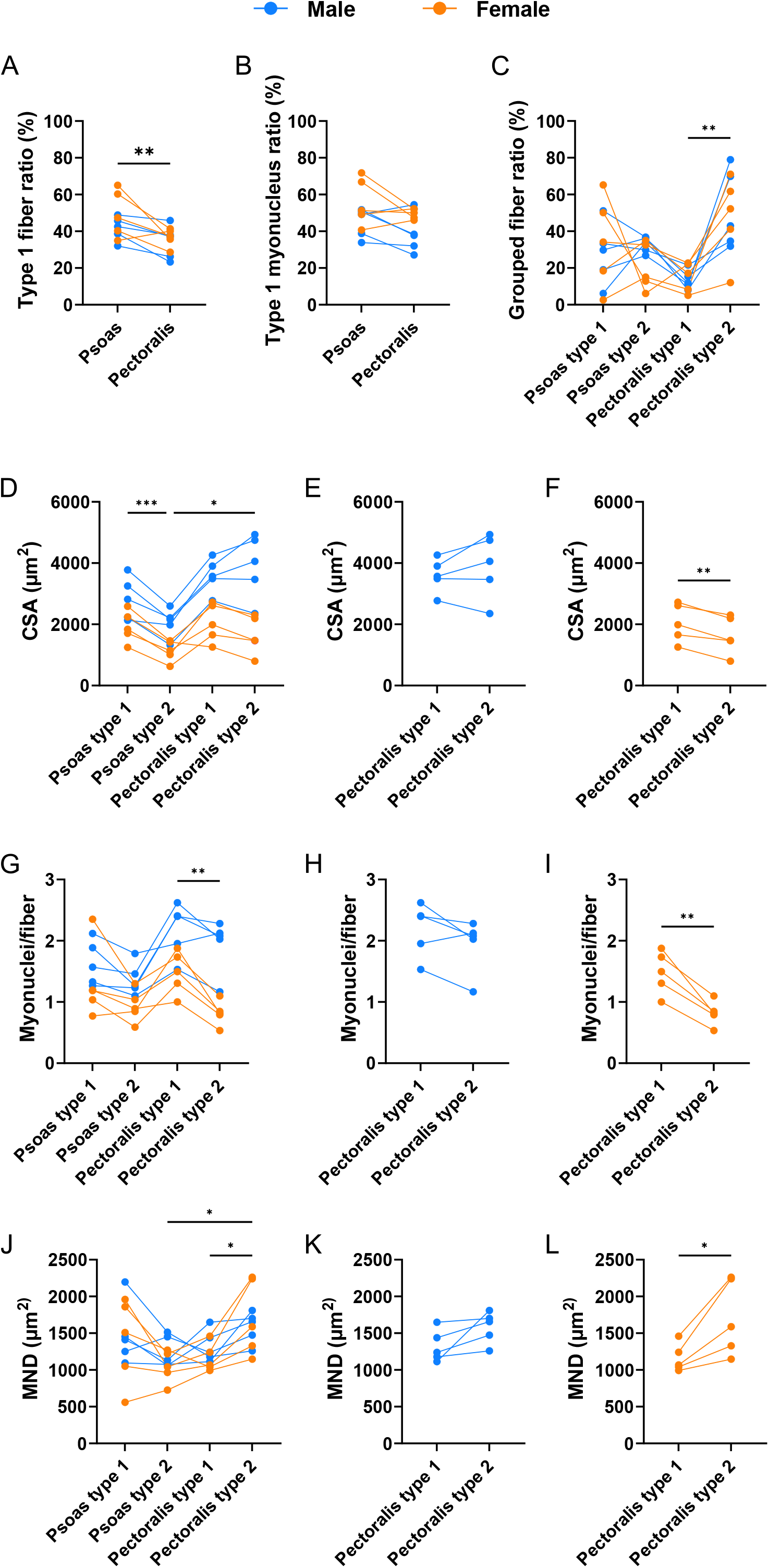
FiNuTyper identifies gender-, muscle- and fiber type-specific characteristics in healthy human skeletal muscle. Fiber- and myonuclear evaluation of healthy skeletal muscle tissue was performed by the FiNuTyper pipeline. Data was collected and pooled from 3, approx. 0.9 × 0.9 µm^2^ image scans of postmortem psoas major and pectoralis major samples of five males (blue data points) and five females (orange data points) between 44-55 years of age. Data points derived from the same subject are visualized with a connecting line. (A) Type 1 fiber and (B) myonucleus ratios of psoas and pectoralis muscle. (C) Grouped fiber ratios of type 1 and type 2 fibers in the psoas and pectoralis muscle. (D) Cross-sectional area (CSA) of type 1 and 2 fibers in the psoas and pectoralis muscle. (E) Analysis of paired male and (F) female CSA data points in pectoralis type 1 and type 2 fibers. (G) Myonucleus per fiber values of type 1 and 2 fibers in the psoas and pectoralis muscle. (H) Analysis of paired male and (I) female data points in pectoralis type 1 and type 2 fibers. (J) Myonuclear domain size (MND) of type 1 and 2 fibers in the psoas and pectoralis muscle. (K) Analysis of paired male and (L) female data points in pectoralis type 1 and type 2 fibers. Statistical analysis was performed between all compared datasets, with one-way ANOVA and Tukey post hoc test (C, D, G, J), paired t-tests (A, B, E, F, I, K, L), or Wilcoxon matched-pairs signed rank test (H), depending on the results of previous normality test. Statistical significance was set to p<0.05 and marked with asterisk (*: p < 0.05; **: p ≤ 0.01; ***: p ≤ 0.001; ****: p ≤ 0.0001; lack of statistical difference was not marked on the plots). Data are presented as mean ± SD.

FiNuTyper allows for the identification of grouped fibers, here defined as myofibers having three or more direct neighbors of the same type ^31^. This value varied most in the functionally dominant psoas type 1 and pectoralis type 2 fibers, with the latter showing a significantly higher grouped fiber ratio compared to type 1 fibers in the same muscle (Fig. 5C, p=0.0025). Some of this difference, however, is due to the distinct fiber type composition of the two studied muscles ^32^. Further mathematical analysis, aiming to minimize the intrinsic bias caused by the unequal abundance of the studied fiber subsets, can benefit from information on fiber type distribution, and other geometric parameters, such as cross-sectional area, readily available for every single fiber after the automated analysis.

The relationship between type 1 and type 2 fiber cross-sectional areas largely depends on the prevalent function of most muscles. Accordingly, in our pooled dataset type 1 fibers displayed a larger CSA than type 2 fibers in the psoas major, supporting its postural role (Fig. 5D, p=0.0001). In the pectoralis with a more dynamic function, we found no significant difference in size between the two fiber types, explained by the type 2 fibers being almost twice as large as their counterparts in the psoas (Fig. 5D, p=0.0184). Males manifested strikingly larger CSA values in all four studied fiber subsets compared to females (Fig. 5D). The relationship between data points collected from the same individual was largely similar over the entire dataset and between genders, except for the CSA of pectoralis type 2 fibers, which seemed to follow distinct trends in males and females. While in males these fibers were as large as pectoralis type 1 fibers (Fig. 5E) and almost twice as big as psoas type 2 fibers (Fig. 5D), in females they showed the opposite pattern, being significantly smaller than the type 1 fibers of the same muscle (Fig. 5F, p=0.0037) and in the same size range as their counterparts in the psoas major (Fig. 5D).

We observed a similar trend to the CSA measurements in the myonuclei per fiber values between psoas type 1 and type 2 fibers (Fig. 5G). On the other hand, we found a pronouncedly higher myonuclear content in type 1 than in type 2 fibers of the pectoralis muscle (Fig. 5G, p=0.0082), in contrast to their similar CSA values. When analyzing the data points of the two genders separately, this difference disappeared in the male cohort (Fig. 5H), while the female pectoralis type 1 and type 2 fibers exhibited a similar relationship as in the pooled dataset (Fig. 5I, p=0.0031). These results corresponded well with the distinct pattern of CSA values seen in the male and female pectoralis muscles (Fig. 5E, F).

Based on the classical concept of the myonuclear domain (MND), the number of myonuclei assigned to single fibers should follow changes in the CSA, however, this assumption has been challenged in recent years ^26,33,34^. We calculated the MND for all four fiber subsets in each subject (Fig. 5J). Pectoralis type 2 fibers on average showed higher MND compared to both psoas type 2 (Fig. 5J, p=0.0289) and pectoralis type 1 fibers (Fig. 5J, p=0.0161). The latter difference, however, was not present separately in the male cohort (Fig. 5K) and was driven by two female subjects (Fig. 5L, p=0.0244), who also had the largest MND in psoas type 1 fibers in the female cohort (Fig. 5J).

All these results indicate gender-specific mechanisms and the relevance of individual-based factors in determining fiber type characteristics of different muscles (Fig. 5A-L). The mean values and standard deviation or all assessed parameters, calculated in the pooled cohort and male and female subjects separately, are displayed in Supplementary Table 2.

### Automated SERCA-based muscle analysis reveals pathological changes in ALS

To test the applicability of the FiNuTyper pipeline to pathological muscle tissue, we analyzed a vastus lateralis muscle biopsy from a female patient with end-stage amyotrophic lateral sclerosis (ALS). First, we examined whether the association between corresponding SERCA and MyHC isoforms, observed in healthy tissue, remains intact in the diseased tissue. We compared MyHC1-MyHC2A-2X- and SERCA1-SERCA2-specific labeling in identical areas of consecutive tissue sections and found almost complete accordance between the signal patterns acquired from the two parallel stainings (Fig. 6A), suggesting that SERCA1 and SERCA2 detection allows for fiber-type discrimination even under pathological conditions.

**Figure 6.**
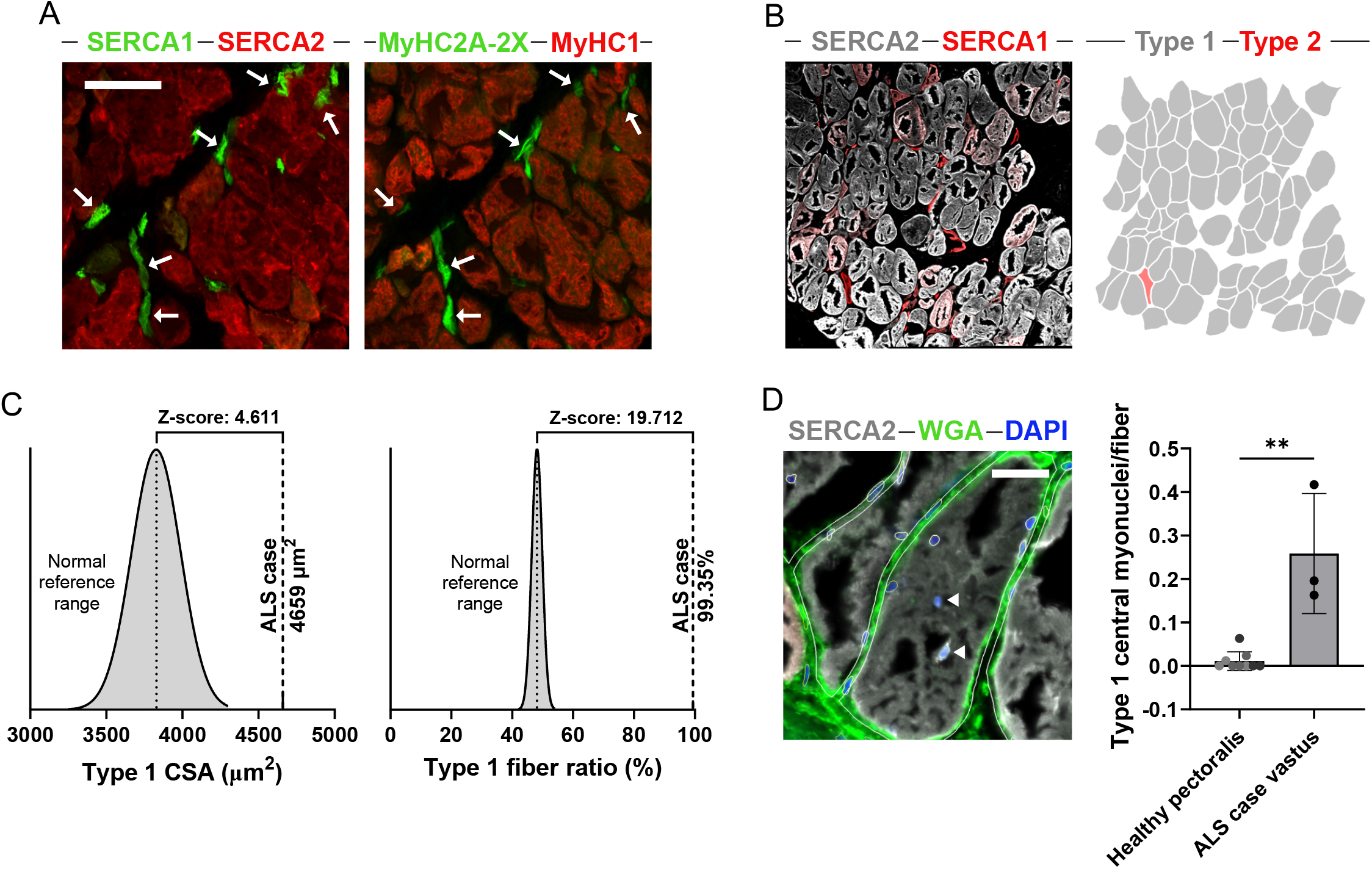
FiNuTyper detects type 2 fiber loss and abundance of central myonuclei in ALS-affected muscle. (A) SERCA1 (green) - SERCA2 (red) (left panel) and MyHC2A-2X (green) - MyHC1 (red) (right panel) labeling provides highly similar staining patterns in consecutive sections of skeletal muscle tissue of an ALS patient. Type 2 fibers identified by SERCA1 or MyHC2A-2X expression are marked by arrows. The scale bar represents 100 µm. (B) Segmentation of type 1 (grey) and type 2 (red) fibers (right panel) based on SERCA2 (white) and SERCA1 (red) immunostaining signal (left panel) in vastus lateralis muscle of an ALS patient. (C) Mean type 1 fiber CSA (µm^2^) and type 1 fiber ratio (%) of the ALS muscle sample, calculated from pooled data of three, approx. 0.9 × 0.9 µm^2^ image scans (n=304 type 1 and n=1 type 2 fibers identified). Comparison to reference values for type 1 fibers of healthy female vastus lateralis tissue ^35^ yields a Z-score of 4,611 for CSA and 19.712 for type 1 fiber ratio, assuming normal distribution of the reference data. (D) Centrally located myonuclei (arrowheads) in type 1 fibers are more frequent in the pathological vastus lateralis sample (black data points) than in the pectoralis major of three healthy female subjects of the same age group (data points in different shades of grey). Three image scans per subject were analyzed and treated as technical replicates. The scale bar represents 40 µm.

The donor suffered from ALS with a bulbar onset for 8 years, and accordingly, we observed severe alterations of the muscle architecture with thickened connective tissue walls between muscle cells and pronounced atrophy and loss of type 2 myofibers (Fig. 6B). As expected, the most severely damaged type 2 fiber remnants were not identified as individual fiber objects by the pipeline, with the few annotated type 2 fibers retaining most resemblance to the normal fiber phenotype (Fig. 6B). We compiled data from three image scans (approx. 0.9 × 0.9 mm2, 306 fibers and 567 myonuclei analyzed in total) of the diseased sample and compared the results to gender-specific reference values of fiber type composition and cross-sectional area, determined by a meta-analysis of 19 independent studies (Fig. 6C) ^35^. The CSA of type 1 fibers in the ALS sample (4659 µm^2^) was above the published reference range (3829 ± 180 µm^2^, mean ± SD) (Fig. 6C), indicating a hypertrophic response in this fiber population. At the same time, the extremely high ratio of type 1 fibers (99.36%), in comparison to the balanced distribution of fiber types according to the healthy reference value (48.1 ± 2.6, mean ± SD), supported our initial finding of substantial type 2 fiber loss in the diseased sample (Fig. 6C). Muscle damage often initiates regenerative processes in the affected tissues with a common manifestation of increased frequency of centrally located myonuclei (Fig. 6D), signaling ongoing myonuclear accretion to the impaired fibers. We compared the number of centrally located myonuclei per type 1 fiber in three technical replicate image scans of the ALS tissue to three healthy, female pectoralis samples in our postmortem dataset and found significantly higher values in the ALS tissue (Fig. 6D, p=0.0103). All these observations are in line with our current understanding of ALS disease progression in human skeletal muscle and confirm that the FiNuTyper pipeline is adept at detecting pathology-related phenomena when analyzing diseased skeletal muscle tissue.

## Discussion

Despite numerous technological advancements of late, immunohistochemical analysis of bioptic samples remains the methodological touchstone of skeletal muscle research. Acknowledging the importance of this approach, a substantial number of automated tools have been presented in recent years (Suppl. Table 1), aiming to increase the speed, accuracy, and extent of the microscopic image evaluation and avoid operator-induced bias, the most obvious limitations of the classical manual quantification. Each of these algorithms assesses biologically relevant myofiber characteristics, however, only a few of them perform simultaneous myonucleus assignment, relying exclusively on positional cues of nuclear objects in relation to the sarcolemma ^16,20^. Challenges of unequivocal myonucleus identification have been pointed out as a major source of controversies around cellular processes involving myonuclear accretion or apoptosis ^5^.

With FiNuTyper, we present the first automated image analysis pipeline utilizing myonuclear markers for skeletal muscle immunohistological evaluation. The only previously reported myonuclear marker, pericentriolar material 1 (PCM-1) ^25^, which is a component of the perinuclear matrix in mature myonuclei, does not allow for direct fiber type assignment ^25^. We used SERCA1- and SERCA2-specific immunostaining to obtain a distinct perinuclear signal on top of the extensive sarcoplasmic reticulum labelling in type 1 and type 2 myofibers, respectively. This approach holds the additional advantage of allowing for simultaneous fiber- and myonucleotyping in the same muscle section.

We performed meticulous validation of the SERCA1- and SERCA2-specific antibodies used in our study. We found no specific SERCA2 labeling in an intensity range similar to that in muscle fibers in any other muscle-resident cell type, although the SERCA2b isoform has been reported to have a ubiquitous expression ^28^.

We performed detailed characterization of MyHC and SERCA isoform expression patterns in various healthy muscle samples and in a pathological tissue. We found a high degree of concordance between the two labeling strategies, even in older individuals, where it has been suggested that the coordination between SERCA and MyHC isoforms might break down ^32^. Notably, in several samples, we detected a subpopulation of SERCA2-positive fibers with an intermediate level of SERCA1 expression (Suppl. Fig. 2A), potentially signaling a wider phenotypical range in the type 1 fiber population and necessitating stringent gating strategies (Suppl. Fig. 2B) when directly comparing the results of MyHC- and SERCA-based staining approaches.

FiNuTyper utilizes a combination of deep learning-based approaches for automated image analysis, allowing for fast, and highly accurate identification and characterization of fiber and myonuclear objects in human skeletal muscle tissue under various experimental conditions. Concerning accuracy performance, we found the quality of tissue and staining in the analyzed image play the largest roles. In our hands, the integrity and conciseness of the fiber border are the most significant contributors to this type of quality difference. Since the quality of tissue is difficult to control for, and there is some confusion regarding how to quantify methodological reliability within the field ^36^, we refrain from directly comparing the accuracy of FiNuTyper to other published automated approaches. The accuracy of myonuclear identification by FiNuTyper, compared to manual evaluation, is still considered good to excellent according to convention ^37^. This is also, however, the feature most affected by poor tissue quality, underlining the technical difficulties in this type of analysis when solely relying on the localization of myonuclei inside of the sarcolemma border. A high level of object recognition fidelity is required for proper fiber shape retention in the small peripheral region where the myonuclei are located, and only a few pixels of error in the identified fiber area might give rise to a neighboring cell’s nucleus appearing within the fiber border and thus being considered a myonucleus. Therefore, including SERCA1 and SERCA2 labeling as a second level of myonucleus identification improves the accuracy of this type of analysis.

We showcased the overall efficiency and robustness of FiNuTyper-based evaluation by analyzing postmortem muscle samples of five male and five female subjects from the same age group (44-55 years). We collected tissue from two different muscles of the deceased donors, the psoas major and the pectoralis major, that are either inaccessible or rarely sampled in studies using biopsies from living subjects. This allowed us to evaluate myocyte- and myonucleus-related parameters at the level of the individual, muscle source (psoas major and pectoralis major), and fiber type (type 1 or type 2).

Moreover, we performed automated analysis on a muscle biopsy of a patient suffering from severe amyotrophic lateral sclerosis. Our analysis demonstrated that FiNuTyper, beyond characteristics of healthy muscle tissue, also successfully identifies features associated with muscle pathology and regenerative processes, such as the loss of fast-twitch motor units and a high frequency of centrally located myonuclei ^38–41^.

The versatility of FiNuTyper lies largely on the Cellprofiler environment ^42,43^, which allows for various adaptations of the original pipeline to answer other relevant research questions in the skeletal muscle field. To demonstrate this, we supply five additional annotated pipelines that build on the same basic principles but produce different outputs or utilize different markers. These include identifying and quantifying the hybrid myofiber and myonuclear populations using the SERCA markers (Fig. 7A), capillarization on a fiber subtype basis using UEA I, satellite cells using PAX7, tissue-resident and infiltrating immune cells using CD45 (Fig. 7B), and fiber-typing based on conventional MyHC markers. Validating these pipelines is beyond the scope of this project, but we supply them, along with example images, as steppingstones for further research (Mendeley data DOI: 10.17632/dfw8r794ph.1).

**Figure 7.**
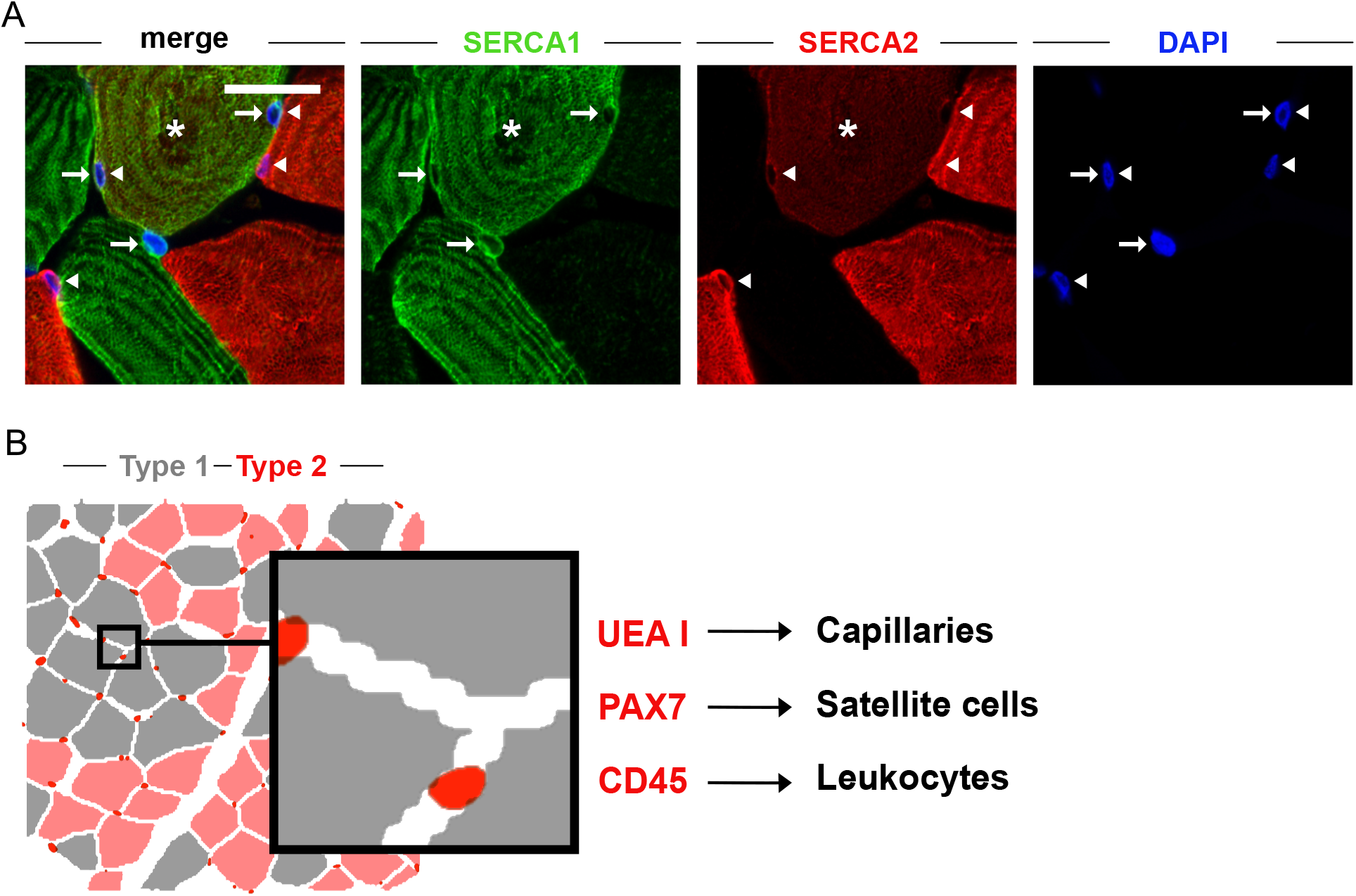
Versatility of FiNuTyper in other experimental settings. (A) Quantification of hybrid myofibers (asterisk) is based on the co-expression of the two SERCA isoforms, which also allows for the identification of hybrid myonuclei with overlapping perinuclear SERCA1 (arrow) and SERCA2 (arrowhead) signals. (B) Schematic overview of modified FiNuTyper applications includes capillary identification based on UEA I labeling, satellite cell identification based on nuclear PAX7 expression, and immune cell identification based on the presence of the CD45 marker.

To conclude, with FiNuTyper we present an automated method for the high-throughput evaluation of the most important aspects of skeletal muscle histology, including simultaneous myofiber and myonucleus type analysis. For this goal, we have introduced and validated SERCA1 and SERCA2 as a novel, type 2- and type 1-specific myofiber and myonuclear markers, respectively. We have shown that our pipeline delivers results in line with human manual evaluation, and is robust enough to discern gender-, muscle origin- and fiber type-specific differences even in a relatively small sample population. The platform can also be successfully applied to pathological muscle samples, and is highly customizable, which we demonstrate by providing five additional pipelines for the analysis of other resident cell types, capillaries, and hybrid myofibers. Together, FiNuTyper facilitates the rapid processing of large image sets and analysis of multiple muscle characteristics, while increasing robustness and reproducibility in image-based skeletal muscle research.

## Methods

### Study subjects, tissue collection, and ethics

Postmortem tissue samples of psoas major and pectoralis major muscles were collected at the Department of Oncology-Pathology of Karolinska Institutet, Sweden, from 21 overall healthy donors during routine autopsies, with the informed consent of relatives (ethical permit number: Dnr 02-418.) Tissue biopsy from the vastus lateralis muscle of an ALS patient was provided by Susanne Petri from Hannover Medical School, Germany (ethical permit number: 6269). Tissue sections of vastus lateralis muscle biopsies, collected from 3 healthy subjects were provided by Carl Johan Sundberg from Karolinska Institute, Sweden (ethical permit number: Dnr 2016/590-31). Gender, age, and sampled muscles of the study subjects are listed in Supplementary Table 3, while a summary of figures and datasets in which images or data from particular study subjects were included is presented in Supplementary Table 4.

### Tissue handling and cryosectioning

The postmortem muscle samples and the tissue from the ALS subject were cut into approximately 5×5×5 mm^3^ segments and then placed in a cryomold filled with Tissue-Tek O.C.T Compound (Sakura), in an orientation allowing for transversal sectioning of the myofibers. The tissue blocks were flash frozen in an isopentane-dry ice slurry, while the fresh bioptic muscle samples were flash frozen in liquid nitrogen-cooled isopentane, and then stored at -80 °C until sectioning. 7 µm (fresh biopsies), 20 µm (ALS tissue) or 40 µm (postmortem tissue) thick sections were cut from the O.C.T-embedded samples in a cryostat with -20 °C chamber temperature and -16 °C blade temperature. The sections were transferred and mounted on glass slides using the CryoJane Tape Transfer system and were either processed immediately or stored at -80°C in a tightly closed container until later use.

### Immunohistochemistry

The slides were quickly warmed up to room temperature and then fixated in 4% formaldehyde-PBS (phosphate-buffered saline) for 20 minutes, followed by washing steps in PBS (3×5 minutes). 250 µl blocking and permeabilization solution (0.1% Triton X-100 and 4% normal goat or donkey serum in PBS), containing different combinations of primary antibodies (anti-SERCA1 (mouse monoclonal IgG1, VE121G9 clone, MA3-912, Thermo Fischer Scientific, 2 µg/ml); anti-SERCA1 (rabbit polyclonal IgG, PA5-78835, Thermo Fischer Scientific, 2.5 µg/ml); anti-SERCA2 (rabbit monoclonal IgG, EPR9392 clone, ab150435, Abcam, 2 µg/ml); anti-MyHC1 (mouse monoclonal IgG2b, BA-D5 clone, DSHB, 1.25 µg/ml); anti-MyHC2A-2X (mouse monoclonal IgG1, SC-71 clone, DSHB, 2.5 µg/ml); anti-lamin A/C (mouse monoclonal IgG2b, 636 clone, sc-7292, Santa Cruz Biotechnology, 1 µg/ml); anti-lamin A/C (goat polyclonal IgG, N-18, sc-6215, 1 µg/ml); anti-PAX7 (mouse monoclonal IgG1, DSHB,2.5 µg/ml); anti-CD45 (mouse monoclonal IgG1, MEM-28 clone, ab8216, 2 µg/ml) and antilaminin (rabbit polyclonal, L9393, 1 µg/ml) was applied on each section, and the slides were then incubated in a humidified chamber at room temperature overnight. The following day, the sections were washed in PBS at room temperature (3×10 minutes), then 250 µl PBS with different combinations of fluorescently labeled goat (A21121, A21127, A21242, A21245 or A31556, Thermo Fischer Scientific, 4 µg/ml) and donkey (711-166-152, 711-546-152, 715-166-150 or 715-546-151, Jackson ImmunoResearch, 1.5 µg/ml) secondary antibodies, Alexa Fluor 488-or 647-conjugated wheat germ agglutinin (WGA) (W11261 or W32466, Thermo Fischer Scientific, 5 µg/ml) and rhodamine-conjugated Ulex Europaeus Agglutinin I (UEA I) (RL-1062, Vector Laboratories, 10 µg/ml) was applied on the slides. The sections were incubated in a dark humidified chamber for 1 h at room temperature, with subsequent washing steps in PBS (3 times 10 minutes). The slides were submersed in PBS containing 0.2 µg/ml 4,6-diamidino-2-phenylindole (DAPI, Thermo Fisher Scientific) for 5 minutes between the second and third washing steps. Finally, glass coverslips were mounted on the sections using ProLong Gold Antifade Reagent (P10144, Thermo Fisher Scientific) and sealed with transparent nail polish after solidification. The stained slides were then stored at 4 °C in dark.

### Confocal microscopy and image processing

Areas of interest of transversally cut muscle regions were selected based on the perceived roundness of myofibers, minimal shift in the fiber border signal between different imaging planes, and a SERCA-labelling pattern consistent with perpendicular fiber orientation. Representative images of the stained skeletal muscle sections were captured by a Carl Zeiss LSM 700 laser scanning microscope with a Zen 2012 Black Edition software, using a Carl Zeiss Plan-Apochromat 20x/0.8, 40x*/*1.3 Oil DIC (UV)VIS-IR and 63x/1.4 Oil DIC objectives, with a 1024×1024 pixel resolution and 4.1 µm, 1.9 µm and 1.1 µm optical thickness, respectively. For the manual and automated image analysis, 3×3 tile scans were captured with 20x magnification (0.156 μm/pixel), 5 µm optical thickness and 2048×2048 pixel resolution, and stitched with 10% overlap. For the determination of parameters including nuclear objects, the stitched scans were split into 9 image frames and processed sequentially, due to memory-related technical constraints. For scan-based analysis, the resolution of the tile scans was instead decreased to a quarter of the original (0.624 μm/pixel) using bicubic interpolation, which saved processing time but had a negligible effect on the accuracy of purely myofiber-related parameters.

Linear adjustments to the brightness and contrast of representative images were performed in Zen 2012 Black Edition and Affinity Designer. For images submitted to manual and automated analysis, care was taken not to oversaturate channels with information-bearing intensity values, as those were used to determine fiber-or myonucleus type. In the WGA or laminin channels, signal continuity and thus improved fiber object recognition was instead prioritized, and thus overexposure was allowed. All images were exported as both separate and merged channels in 8-bit png file format and submitted to manual or automated image analysis.

### Image analysis and data generation

The FiNuTyper workflow combines two open-source software with an in-house-developed Cellprofiler ^42^ pipeline. Firstly, nuclear object recognition from the DAPI signal was performed using the image style transfer-based neural network NucleAIzer ^30^, on image frames where average nuclear diameter was set to 30 pixels (0.156 μm/pixel). Then, fiber objects were identified from the myofiber border marker WGA or laminin signal by Cellpose ^29^, on both image frames and stitched scans, with an object diameter of 300 pixels (0.156 μm/pixel, frame-based analysis) or 75 pixels (0.624 μm/pixel, scan-based analysis), with a cytoplasm model and the model’s predefined thresholds. The generated nucleus- and fiber object-masked images, along with a merged and separate channel images for SERCA1, SERCA2, WGA and DAPI, were used as input in the Cellprofiler ^42^ pipeline for the frame-based analysis. No nuclear identification was performed as part of the scan-based analysis and thus no nucleus object-masked image was used in it. All parameters involving myonuclear values (type 1 myonuclear ratio, myonuclei per fiber, central myonuclei per fiber, myonuclear domain) were calculated from the high-resolution images with 0.156 μm/pixel scaling, while purely fiber-related parameters (type 1 fiber ratio, cross-sectional area, grouped fiber ratio) were generated from the whole scans with decreased resolution with 0.624 μm/pixel scaling, allowing for the analysis of a higher number of fibers in the same area of interest.

For fiber and myonucleus type determination, the intensities of the SERCA1 and SERCA2 signals were measured within the fiber and nucleus objects and quantified as average pixel intensity values (mean fiber or nucleus intensity). Fibers located on or close to the image borders were not identified by the pipeline, and fiber objects without measurable SERCA1 or SERCA2 signal were categorized as artefacts and excluded from subsequent analysis. After observing numerous (mainly type 2) fibers of uncommon shapes in our sections, we opted to completely exclude a roundness filter, commonly used in other automated pipelines, to avoid introducing type-specific technical bias in our data collection. The SERCA1 and SERCA2 intensity levels, used for fiber- and nucleotype gating, were calibrated for every experimental setting, using the histogram and density plot features of Cellprofiler Analyst ^43^. For myonucleus type analysis, adjacency (adjacent object pixels or overlap) to a fiber object of the similar type was also considered. Based on the more distinct profile of SERCA2-positive fibers and myonuclei, fiber and nuclear objects co-expressing SERCA1 and SERCA2 were annotated as type 1.

Data used for subsequent analysis were extracted from the raw data file generated by the Cellprofiler ^42^ pipeline and compiled per individual or per image scan (technical replicates). Individual numbers assigned to myofiber objects in the output dataset allowed for their unequivocal identification in the original images, making post-hoc manual correction possible. Primary derived values (number of fibers and myonuclei, fiber size, number of central myonuclei and number of grouped fibers) were compiled and averaged in a type-specific manner and were used to calculate secondary derived values (type 1 fiber percentage, type 1 myonucleus percentage, myonucleus per fiber, myonuclear domain, grouped fiber percentage and grouped fiber size compared to average fiber size). These values were then used to characterize distinct fiber populations of the study subjects.

In the five additional, annotated pipelines, the following minor modifications were made: fiber type annotation was performed exclusively based on the presence or absence of SERCA2 signal and the SERCA1 channel was replaced by UEA I (capillary identification), CD45 (leukocyte identification) or PAX7 (satellite cell identification). For capillary identification, nuclear objects were not considered, but UEA I-positive objects were annotated as capillaries using NucleAIzer (average object diameter set to 30 pixels). For leukocyte and satellite cell annotation, an overlap between DAPI and CD45 or Pax7 signals were considered. For a classical fiber typing pipeline, the SERCA1 channel was exchanged to MyHC2A-2X and the SERCA2 to MyHC1 and the WGA to laminin. Finally, for the quantification of hybrid fibers and myonuclei, parallel gating strategy for SERCA1 and SERCA2 and identification of double positive objects was implemented.

The in-house-developed Cellprofiler module of the FiNuTyper pipeline is provided in Supplementary File 1 (scan-based analysis) and Supplementary File 2 (frame-based analysis), while technical notes and detailed instructions on how to run FiNuTyper is presented in Supplementary File 3 (Mendeley data DOI: 10.17632/dfw8r794ph.1).

### Validation of MyHC and SERCA isoform co-expression

Overlap between the different MyHC and SERCA isoforms was evaluated in three male subjects of 25, 45, and 73 years of age, in immunostainings combining MyHC1-, MyHC2A-2X- and SERCA1-or SERCA2-specific antibodies with fluorescently conjugated WGA. Three, approx. 0.9 mm × 0.9 mm^2^ image scans (3×3 image frames with 10% overlap) were captured from each individual and were submitted to automated image analysis. Per scan, 213.78 ± 25.47 fibers for SERCA1 and 234.78 ± 20.41 fibers for SERCA2 validation were identified (mean ± SD). Mean fiber intensities were measured in all three relevant channels. After manual exclusion of incorrectly outlined fiber objects, mean fiber intensity in the SERCA channel, measured in individual fiber objects, was plotted against the mean fiber intensity in the corresponding MyHC channel (SERCA1–MyHC2A-2X; SERCA2–MyHC1). Quadrant gates were defined by the lowermost intensity values in the clearly double-positive population, outlining true positive (TP; SERCA^+^-MyHC^+^), true negative (TN; SERCA^-^-MyHC^-^), false positive (SERCA^+^-MyHC^-^) and false negative (SERCA^-^-MyHC^+^) fiber populations in each analyzed image scan. Sensitivity (TP/(TP+FN)), specificity (TN/(TN+FP)), positive predictive value (TP/(TP+FP)) and negative predictive value (TN/(TN+FN)) were calculated for each scan, and then used to generate mean values for each subject separately. Final accuracy values were calculated as an average for the three subjects, displayed as mean ± SD.

### Validation of fiber and myonucleus identification and CSA measurements

Fiber and myonucleus identification in postmortem psoas major and pectoralis major sections was evaluated by 2 independent operators, blind to the generated count, age, and gender of the subjects, in a 1/3 overlapping fashion (n=57 randomly selected, approx. 0.3 × 0.3 mm^2^ image frames from ten subjects containing 880 fibers and 1179 myonuclei in total). Operator 1 validated 25 such frames, operator 2 validated 23 such frames, and 9 such frames were validated by both operators, the results of which were averaged for subsequent evaluation. Similar analysis of frozen sections of fresh vastus lateralis muscle biopsies (n=9 randomly selected approx. 0.3 × 0.3 mm^2^ image frames from three subjects containing 102 fibers and 304 myonuclei in total) was performed by a single operator. Validation of cross-sectional area measurements was performed separately in type 1 and type 2 fibers by a single, similarly blinded operator, both in postmortem (n=35 randomly selected, approx. 0.3 × 0.3 mm^2^ image frames from ten subjects containing 554 fibers in total) and fresh bioptic samples (n=9 randomly selected, approx. 0.3 × 0.3 mm^2^ image frames from three subjects containing 102 fibers in total), where mean CSA per image frame values were calculated and displayed. The level of agreement between the manual and automated analyses was determined by calculating intraclass correlation coefficient (ICC) with 95% confidence interval, and Bland-Altman analysis. Automated cross-sectional area measurement was also validated against manual quantification on the individual fiber level in a single image scan (9 stitched image frames, approx. 0.9 × 0.9 mm^2^, n=243 fibers identified automatically; n=254 fibers evaluated manually).

### Statistics and data presentation

Intraclass correlation coefficient (ICC) analysis (two-way random effects, absolute agreement, single measures) was performed in IBM SPSS statistics 27 to assess comparability between FiNuTyper-based and manual evaluation. Bland-Altman analysis for the validation dataset, paired data and group comparisons and visualization were performed in GraphPad Prism 9.3.1.

Normal distribution of grouped datasets was analyzed by Shapiro-Wilk and Kolmogorov-Smirnov tests. Differences between groups were analyzed either with one-way ANOVA and Tukey’s post-hoc test, or in case of two groups, paired t-tests, Wilcoxon matched-pairs signed rank test, or Mann Whitney U test, depending on analyzing paired or unpaired data points and the result of the previous normality test. Statistical significance was set to p<0.05 and marked with asterisk (*: p<0.05; **: p≤0.01; ***: p≤0.001; ****: p≤0.0001). For the validation analysis, per-image values were calculated and compared in all assessed parameters. For the analysis of the healthy subject cohort, generated values were pooled on a per-subject basis. Central myonuclei per fiber values were compared on a scan-basis between 3 healthy and one pathological muscle samples. Z-score was calculated from the Z=(x-µ)/σ formula. All data are presented as mean ± SD.

## Data and code availability

Data and code used in this study are available on Mendeley data (DOI: 10.17632/dfw8r794ph.1) and will be made public upon publication.

## Author contributions

A.L. designed the study, devised FiNuTyper, performed and evaluated experiments, analyzed data, prepared figures, and wrote the manuscript. E.L. designed the study, performed and evaluated experiments, analyzed data, prepared figures, wrote and revised the manuscript, and supervised the project. N.S.H. performed and evaluated experiments. E.B.E., S.M.R., M.A.C., K.A., H.D., S.P., and C.J.S. collected tissue samples for the study. O.B. designed the study, wrote and revised the manuscript, supervised the project, and provided funding. All authors approved the final version of the manuscript.

## Acknowledgements

O.B. was supported by the Center for Regenerative Therapies Dresden, the Karolinska Institutet, the Swedish Research Council, the Ragnar Söderberg Foundation, the Åke Wiberg Foundation, and the LeDucq foundation. S.M.R was supported by a doctoral grant from Karolinska Institutet. C.J.S., E.B.E. and M.A.C. were supported by grants from the Swedish Research Council and the Swedish Research Council for Sport Science. We would like to thank Marion Baniol for her help with handling the postmortem muscle samples and discussions about the project, and Paula Heinke and Wouter Derks for their insightful comments on the manuscript.

## Figure legends

**Supplementary figure 1. Perinuclear SERCA1 and SERCA2 labeling is specific for myonuclei**. (A) SERCA1 (arrows) and SERCA2 (arrowheads) show a distinct labelling pattern around nuclei inside of skeletal muscle fibers, but not around non-myonuclei (drop). The scale bar represents 20 µm. (B) No SERCA1-or SERCA2-specific staining can be observed in vessel walls or larger connective tissue segments labeled by WGA, in skeletal muscle sections. The scale bar represents 100 µm. (C) Satellite cells, identified by nuclear PAX7 labeling, do not display SERCA1-or SERCA2-specific nuclear or perinuclear signals. The scale bar represents 10 µm.

**Supplementary figure 2. A subpopulation of type 1 fibers expresses intermediate levels of SERCA1**. (A) Combined SERCA1-SERCA2 immunostaining detects a subpopulation of SERCA2-positive fibers with weak SERCA1 signal (asterisk) in a muscle sample with no hybrid fibers co-labelled by MyHC1 and MyHC2A-2X. The scale bar represents 200 µm. (B) Automated measurements of mean fiber intensities in the MyHC1, MyHC2A-2X, SERCA1, and SERCA2 channels identify a small group of type 1 fibers with intermediate SERCA1 expression (orange data points), present both in subject #1 with virtually no, and subject #2 with a higher ratio of real hybrid fibers (yellow data points), defined by co-expression of MyHC1 and MyHC2A-2X. By using stringent dual thresholds for SERCA 1 and SERCA2 intensities, hybrid fibers appear in similar proportions to those in MyHC-stained samples from the same individual. N=234 (upper left panel), n=241 (upper right panel), n=269 (lower left panel), and n=224 (lower right panel) fibers analyzed and displayed.

**Supplementary figure 3. Bland-Altman analysis of validation dataset**. Difference between manual and FiNuTyper-based analysis of randomly selected image frames, displayed as ratio (manual/automated) vs. average, with 95% limits of agreement (dotted lines). Agreement between the two approaches in the (A) number of identified fibers, (B) myonuclei, (C) and cross-sectional area of type 1 and (D) type 2 fibers was assessed separately in fresh biopsies (left panels) and postmortem muscle (right panels). A detailed evaluation of the validation dataset is presented in Figure 4.

**Supplementary Table 1. Overview of features in selected automated muscle analysis approaches**. ^11,13,14,16,19,22–24^

**Supplementary Table 2. Mean values and standard deviation of fiber- and myonucleus-related parameters in the healthy subject cohort, determined by FiNuTyper**.

**Supplementary Table 3. Information on subjects included in this study**.

**Supplementary Table 4. List of subjects used for different figures and datasets**.

