## Supplemental Figures + Tables for "FiNuTyper: an automated deep learning-based platform for simultaneous fiber and nucleus type analysis in human skeletal muscle"

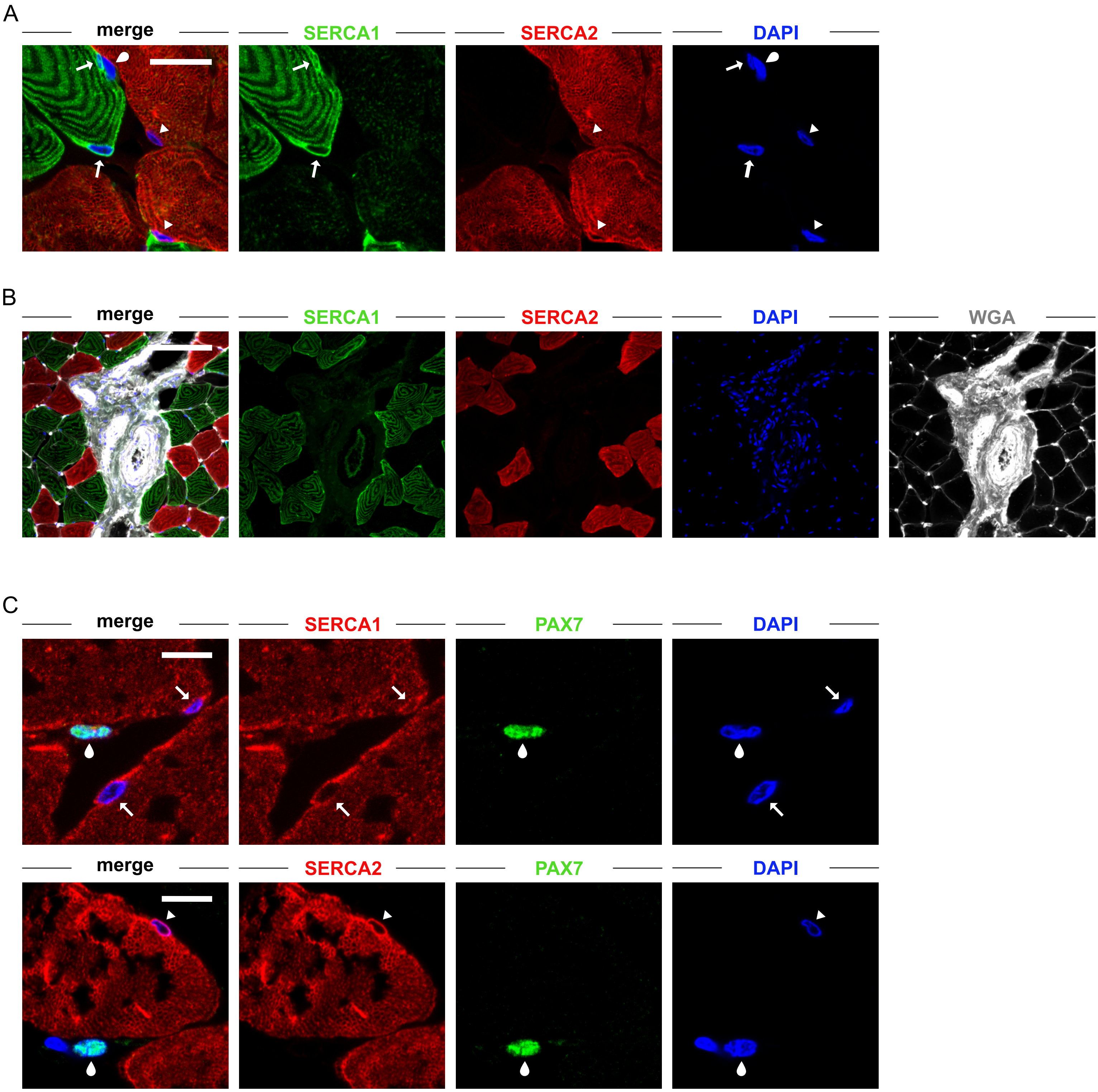


**Supplementary Figure 1**

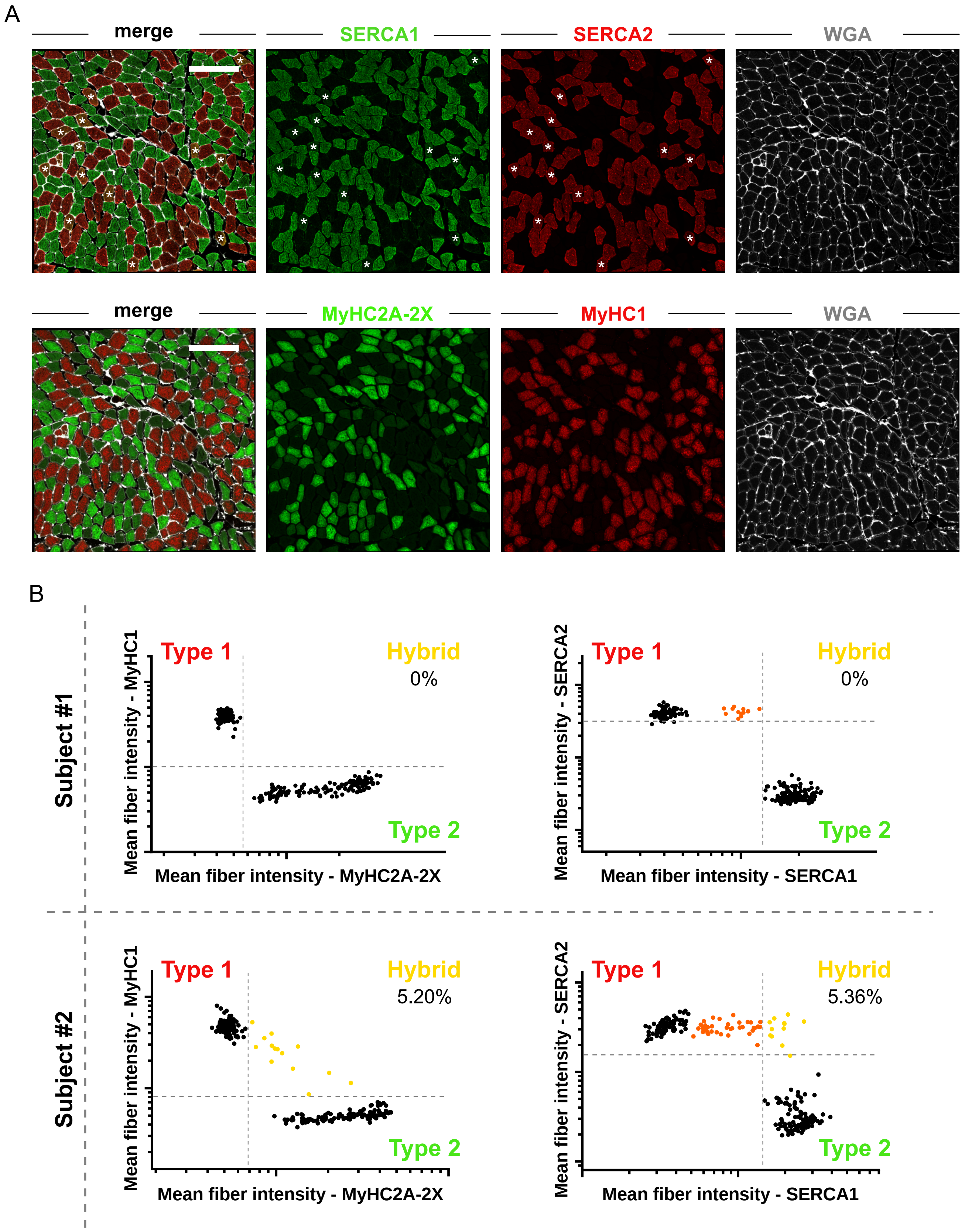


**Supplementary Figure 2**

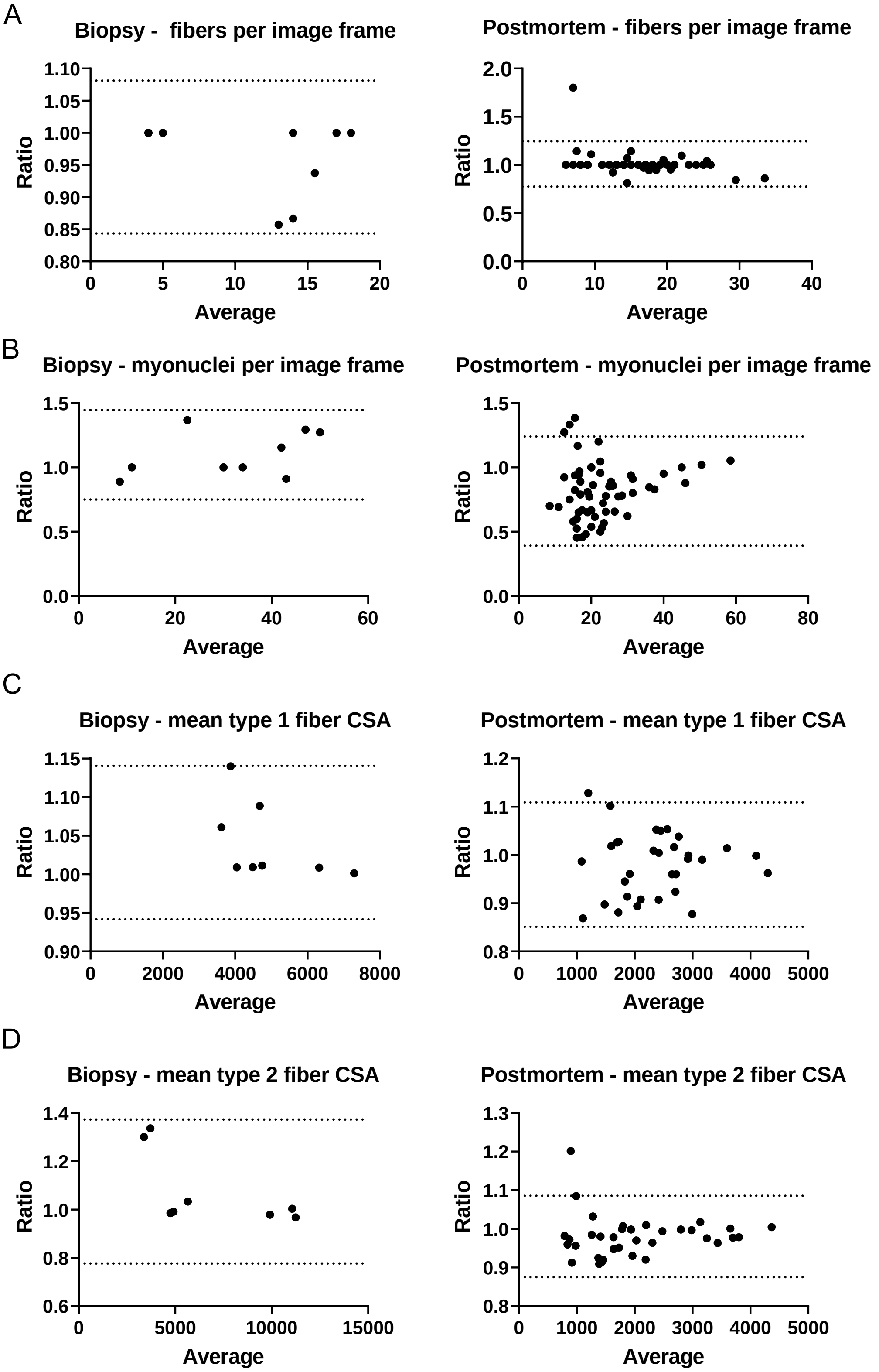


**Supplementary Figure 3**

| Feature | FiNuTyper | Myovision 2.0 | Cellpose application | Myosight | Myosoft | Muscle2view | MuscleJ | Kim et al. | Klemencic et al. |
|---|---|---|---|---|---|---|---|---|---|
| **Year of publication** | 2022 | 2022 | 2021 | 2020 | 2020 | 2019 | 2018 | 2007 | 1998 |
| **Fiber identification** | + | + | + | + | + | + | + | + | + |
| **Fiber typing** | + | + | - | + | + | + | + | - | - |
| **Fiber size determination** | + | + | + | + | + | + | + | + | + |
| **Fiber grouping analysis** | + | - | - | - | - | - | - | - | - |
| **Nucleus identification** | + | + | - | + | - | + | + | - | - |
| **Myonucleus identification** | + | ~ | - | ~ | - | ~ | ~ | - | - |
| **Nucleotyping** | + | ~ | - | ~ | - | ~ | ~ | - | - |
| **Central myonucleus identification** | + | - | - | + | - | - | + | - | - |
| **Central myonucleus typing** | + | - | - | + | - | - | + | - | - |
| **Myonuclei per fiber determination** | + | + | - | + | - | - | + | - | - |
| **Myonuclear domain determination** | + | + | - | + | - | - | + | - | - |
| **Hybrid fiber typing** | (+) | - | - | - | + | - | - | - | - |
| **Hybrid nucleotyping** | (+) | - | - | - | - | - | - | - | - |
| **Satellite cell identification** | (+) | - | - | - | - | - | + | - | - |
| **Immune cell identification** | (+) | - | - | - | - | - | - | - | - |
| **Vessel analysis** | (+) | - | - | - | - | + | + | - | - |
| **Platform** | Cellprofiler + Standalone | Standalone | Standalone | Fiji | Fiji | Cellprofiler | Fiji | Automatic standalone | Semi-automatic standalone |

+ yes
- no
~ without myonuclear marker
(+) modified pipeline provided

**Supplementary Table 1**

| | Pooled (n=10) | | | |
|---|---|---|---|---|
| | Psoas major | | Pectoralis major | |
| Type 1 fiber ratio (%) | 45.58 ± 10.5 | | 35.36 ± 7.132 | |
| Type 1 myonucleus ratio (%) | 50.39 ± 11.71 | | 43.75 ± 9.392 | |
| | Type 1 | Type 2 | Type 1 | Type 2 |
| Grouped fiber ratio (%) | 30.98 ± 20.22 | 26.51 ± 11.08 | 14.34 ± 6.435 | 49.63 ± 20.96 |
| Cross-sectional area (μm²) | 2383 ± 751.3 | 1595 ± 620.3 | 2823 ± 989.5 | 2779 ± 1443 |
| Myonuclei/fiber | 1.471 ± 0.5026 | 1.151 ± 0.3395 | 1.833 ± 0.5238 | 1.375 ± 0.672 |
| Myonuclear domain (μm²) | 1434 ± 483.5 | 1146 ± 231.3 | 1242 ± 212 | 1647 ± 378.9 |

| | Male (n=5) | | | |
|---|---|---|---|---|
| | Psoas major | | Pectoralis major | |
| Type 1 fiber ratio (%) | 41.56 ± 6.538 | | 34.08 ± 9.235 | |
| Type 1 myonucleus ratio (%) | 44.82 ± 7.987 | | 37.95 ± 10.33 | |
| | Type 1 | Type 2 | Type 1 | Type 2 |
| Grouped fiber ratio (%) | 27.88 ± 16.77 | 32.71 ± 4.31 | 13.58 ± 5.229 | 51.65 ± 21.43 |
| Cross-sectional area (μm²) | 2842 ± 692.3 | 2056 ± 466 | 3599 ± 555.2 | 3912 ± 1049 |
| Myonuclei/fiber | 1.634 ± 0.3637 | 1.37 ± 0.2686 | 2.182 ± 0.4381 | 1.935 ± 0.4406 |
| Myonuclear domain (μm²) | 1481 ± 424.4 | 1247 ± 217.7 | 1324 ± 219 | 1580 ± 215.8 |

| | Female (n=5) | | | |
|---|---|---|---|---|
| | Psoas major | | Pectoralis major | |
| Type 1 fiber ratio (%) | 49.6 ±12.84 | | 36.64 ± 5.006 | |
| Type 1 myonucleus ratio (%) | 55.97 ± 12.93 | | 49.55 ± 2.786 | |
| | Type 1 | Type 2 | Type 1 | Type 2 |
| Grouped fiber ratio (%) | 34.09 ± 24.79 | 20.32 ± 12.72 | 15.1 ± 8.024 | 47.62 ± 22.78 |
| Cross-sectional area (μm²) | 1924 ± 513.8 | 1133 ± 340.2 | 2047 ± 623.3 | 1647 ± 615.9 |
| Myonuclei/fiber | 1.307 ± 0.6078 | 0.9329 ± 0.2608 | 1.483 ± 0.3463 | 0.8163 ± 0.2003 |
| Myonuclear domain (μm²) | 1388 ± 583.4 | 1045 ± 217.8 | 1160 ± 190.9 | 1713 ± 515.2 |

n=10 for pooled samples, n=5 for males and n=5 for females.
All data are displayed as mean ± SD.

**Supplementary Table 2**

| Subject code | Gender | Age (years) | Sampled muscles | Sample type | Known muscle pathology |
|---|---|---|---|---|---|
| **pm #1** | male | 55 | psoas major, pectoralis major | postmortem | none |
| **pm #2** | male | 54 | psoas major, pectoralis major | postmortem | none |
| **pm #3** | male | 55 | psoas major, pectoralis major | postmortem | none |
| **pm #4** | male | 52 | psoas major, pectoralis major | postmortem | none |
| **pm #5** | male | 44 | psoas major, pectoralis major | postmortem | none |
| **pm #6** | female | 55 | psoas major, pectoralis major | postmortem | none |
| **pm #7** | female | 46 | psoas major, pectoralis major | postmortem | none |
| **pm #8** | female | 49 | psoas major, pectoralis major | postmortem | none |
| **pm #9** | female | 55 | psoas major, pectoralis major | postmortem | none |
| **pm #10** | female | 45 | psoas major, pectoralis major | postmortem | none |
| **pm #11** | male | 45 | psoas major, pectoralis major | postmortem | none |
| **pm #12** | male | 25 | psoas major, pectoralis major | postmortem | none |
| **pm #13** | male | 73 | psoas major, pectoralis major | postmortem | none |
| **pm #14** | male | 50 | psoas major, pectoralis major | postmortem | none |
| **pm #15** | male | 26 | psoas major, pectoralis major | postmortem | none |
| **pm #16** | female | 15 | psoas major, pectoralis major | postmortem | none |
| **pm #17** | male | 37 | psoas major, pectoralis major | postmortem | none |
| **pm #18** | male | 52 | psoas major, pectoralis major | postmortem | none |
| **pm #19** | male | 53 | psoas major, pectoralis major | postmortem | none |
| **pm #20** | male | 50 | psoas major, pectoralis major | postmortem | none |
| **pm #21** | male | 16 | psoas major | postmortem | none |
| **b #1** | male | 52 | vastus lateralis | fresh biopsy | none |
| **b #2** | male | 42 | vastus lateralis | fresh biopsy | none |
| **b #3** | male | 49 | vastus lateralis | fresh biopsy | none |
| **als #1** | female | 50 | vastus lateralis | postmortem | none |

**Supplementary Table 3**

| Subject code | Used in figures | Used in datasets |
|---|---|---|
| pm #1 | Fig. 5A-E, G, H, J, K | Suppl. Data 3 |
| pm #2 | Fig. 5A-E, G, H, J, K | Suppl. Data 3 |
| pm #3 | Fig. 4A-D; Fig. 5A-E, G, H, J, K; Suppl. Fig. 2A, B; Suppl. Fig. 3A-D | Suppl. Data 1, Suppl. Data 2, Suppl. Data 3 |
| pm #4 | Fig. 4A-D; Fig. 5A-E, G, H, J, K; Fig. 6C; Suppl. Fig. 3A-D | Suppl. Data 2, Suppl. Data 3 |
| pm #5 | Fig. 5A-E, G, H, J, K | Suppl. Data 3 |
| pm #6 | Fig. 4A-D; Fig. 5A-D, F, G, I, J, L; Fig. 6C; Suppl. Fig. 3A-D | Suppl. Data 2, Suppl. Data 3 |
| pm #7 | Fig. 4A, B; Fig. 5A-D, F, G, I, J, L; Suppl. Fig. 3A, B | Suppl. Data 2, Suppl. Data 3 |
| pm #8 | Fig. 4A-D; Fig. 5A-D, F, G, I, J, L; Fig. 6C; Suppl. Fig. 3A-D | Suppl. Data 2, Suppl. Data 3, Suppl. Data 4 |
| pm #9 | Fig. 1B; Fig. 4A-D; Fig. 5A-D, F, G, I, J, L; Fig. 6C; Suppl. Fig. 3A-D | Suppl. Data 2, Suppl. Data 3, Suppl. Data 4 |
| pm #10 | Fig. 4A-D; Fig. 5A-D, F, G, I, J, L; Fig. 6C; Suppl. Fig. 3A-D | Suppl. Data 2, Suppl. Data 3, Suppl. Data 4 |
| pm #11 | Fig. 3A-F | Suppl. Data 1 |
| pm #12 | Fig. 2B; Fig. 3F | Suppl. Data 1 |
| pm #13 | Fig. 3F | Suppl. Data 1 |
| pm #14 | Fig. 2A; Suppl. Fig. 2B | Suppl. Data 1 |
| pm #15 | Fig. 1A; Fig. 7A; Suppl. Fig. 1A, B | |
| pm #16 | Fig. 7B | |
| pm #17 | Suppl. Fig. 1C | |
| pm #18 | Fig. 4A-D; Suppl. Fig. 3A-D | Suppl. Data 2 |
| pm #19 | Fig. 4A-D; Suppl. Fig. 3A-D | Suppl. Data 2 |
| pm #20 | Fig. 4A-D; Suppl. Fig. 3A-D | Suppl. Data 2 |
| pm #21 | Fig. 4E | Suppl. Data 2 |
| b #1 | Fig. 4A-D; Suppl. Fig. 3A-D | Suppl. Data 2 |
| b #2 | Fig. 4A-D; Suppl. Fig. 3A-D | Suppl. Data 2 |
| b #3 | Fig. 4A-D; Suppl. Fig. 3A-D | Suppl. Data 2 |
| als #1 | Suppl. Fig. 6A-D | Suppl. Data 2 |

**Supplementary Table 4**
